# A small set of morphological metrics describes wing-planform plasticity: a correlational analysis in a population of the European grapevine moth *Lobesia botrana* (Lepidoptera: Tortricidae)

**DOI:** 10.64898/2026.11.26.765887

**Authors:** Oleg V. Orlov, Evgeniya G. Yurchenko

## Abstract

The insect wing is a complex morpho-functional organ, and it remains unclear whether a set of well-defined metrics can fully capture wing-planform variation. To address this question, we digitised 339 male forewings of the European grapevine moth, *Lobesia botrana* (Lepidoptera: Tortricidae), an economically important grapevine pest. Specimens came from two non-overlapping seasonal flight generations. We applied two complementary shape-analysis methods — Generalised Procrustes Analysis (GPA) and Elliptic Fourier Analysis (EFA) — and correlated their principal components with four flight-related morphological metrics: aspect ratio (AR), a proxy for the second moment of wing area (MA), sweep angle (SA), and apex taper (Taper). For both methods, each of the first four principal components was most strongly correlated with its corresponding metric; together, these components represented 92.3% (GPA) and 95.7% (EFA) of the total variation of the shape. Both methods yielded concordant results, indicating that a small set of interpretable metrics captures the major axes of wing-planform variation. Generations differed significantly in mean MA, AR, and SA, and in Taper variance, consistent with seasonal phenotypic plasticity. These characters can be calculated from a small number of landmarks and therefore can serve as wing-shape markers. The analysis pipeline is implemented in the open-source MorphoGet software. This protocol provides an interpretable summary of wing-planform variation, applicable to flight-related morphology, ecology and evolution, pest monitoring, and other fields.

## Introduction

Flight is one of the key evolutionary innovations of insects and a major driver of their ecological success and species diversity [1, 2]. Active flight facilitates migration, resource acquisition, predator avoidance, and colonisation of diverse habitats [3–5]. In this context, the wing serves not merely as a locomotor organ but as a complex morphofunctional system linking organismal structure, behaviour, and environmental conditions [4, 6–8]. Wing shape is therefore an important morphological trait that reflects ecological specialisation and functional capability [9–11].

Most studies of wing morphology have addressed taxonomic differences, phylogenetic relationships, and flight type [7,12–14]. Intraspecific variety also deserves attention: it provides a framework for investigating developmental pathways and morphs within a species [15] and, in applied contexts, may reflect population-level adaptive potential, local conditions, individual physiological state, and environmental stressors [16–19]. Wing shape can thus capture not only major evolutionary distinctions but also finer microevolutionary and ecological processes within species [7, 19]. When particular shape characteristics consistently distinguish intraspecific groups or correlate with environmental conditions, they can serve as rapid morphological indicators [18, 20, 21]. This approach may support studies of flight-related morphology, evolutionary biology, and ecological monitoring, as well as habitat quality assessment and the integration of morphological metrics with genetic and other data. Phenotypic plasticity also provides a model for understanding how genetic factors and environmental signals shape ontogenetic trajectories and ecomorphs [15, 22–24].

Studies of wing shape have traditionally relied on linear morphometry, venation analysis, and measurements of the wing planform [25, 26]. Currently, numerous flight-related metrics have been proposed. Aspect ratio (AR) and the second moment of wing area (MA) are established, whereas sweep angle (SA) and apex taper (Taper) are less widely used; all metrics are commonly calculated from simplified geometric relationships and approximations (e.g., length, area, angles) [27], and each captures a distinct aspect of wing geometry: AR — elongation, MA — spanwise area distribution, SA — longitudinal-axis inclination, and Taper — apex shape [7,9,28–43]. Here, we focus on these four as a small, interpretable selection from a much larger pool. In parallel, geometric morphometric methods have enabled researchers to analyse shape as an integrated whole using homologous landmarks and Procrustes analysis (GPA) [44, 45], contour-based methods [46], and Elliptic Fourier Analysis (EFA) [47].

Despite this methodological expansion, the morphological variation of insect wings remains incompletely characterised. Geometric morphometric studies generally focus on multivariate disparity (group differences), and the resulting shapes have limitations, as they are often meaningful only within a given analysis [48]. These studies also rely predominantly on venation, because vein junctions are homologous and readily reproducible [23, 49]. The wing planform, by contrast, shapes the aerodynamic profile and mediates the wing’s interaction with airflow during the stroke, yet planform analyses are often hampered by the scarcity of clear, homologous landmarks on the wing surface [24,25,50]. Research has consequently followed two broad approaches — identifying and exploring functional flight-related metrics [7, 9, 19, 26, 28] and analysing integrated shape and disparity with geometric morphometric methods [23] — yet no unified framework exists for describing wing-planform variation consistently across studies. This has led to a proliferation of studies that either search for parameters via heuristic search or identify study-specific shapes. Moreover, no study has yet described wing-shape variation using a limited set of interpretable metrics, and the question of their sufficiency remains open.

Charles P. Ellington’s classical work found strong correlations among different measures of area distribution across flying animals, suggesting that this aspect of wing variation may be summarised by a single parameter [9]. These observations motivate our hypothesis that a limited set of simple metrics describes the principal intraspecific variation in wing planform. However, we did not address size-related variation, including wing and habitus size or wing loading (i.e., the allometry of body weight relative to wing area). In addition, we propose the metrics and their proxies as practical markers for future research.

## Methods

### Study organism and data collection

We studied male forewings of the European grapevine moth, *Lobesia botrana* (Denis & Schiffermüller, 1775) (Lepidoptera: Tortricidae), an economically important grapevine pest. This small Mediterranean moth has a wingspan of 10–13 mm; its larvae feed on flowers and berries, damaging the reproductive organs of the vine and spreading rot, and under favourable conditions the moth can substantially reduce grape yields within infested areas. In most of the world’s viticultural regions it is an invasive or quarantine pest [51–53].

Males were collected during the 2022 season using delta-type pheromone traps baited with (E,Z)-7,9-dodecadienyl acetate at a commercial vineyard on the Taman Peninsula (Western Ciscaucasia, Krasnodar Krai, Russia). Pheromone-trap catches were used to monitor flight activity. We collected adults from two seasonally non-overlapping generations: 112 spring-generation specimens in May and 227 summer-generation specimens in July.

One wing was removed from each individual, brushed clean of scales, and photographed flat beneath a microscope coverslip. Digitisation was performed using TPSUtil and TPSDig2 [54]. A bounding curve was drawn along the distal portion of the wing, from the apex of the pterostigma (costal region) to the point where the anal vein A(1+2) meets the wing margin at the tornus. Fifteen points, including the two endpoints (landmarks 2–16) were placed at equal intervals along the curve. The two endpoints— the pterostigma apex (landmark 2) and the tornus (landmark 16)— anchored the curve. An additional landmark was placed at the base of the radial stem (radius), marking the wing base (landmark 1). The digitisation scheme is shown in Figure 1. Data were recorded in TPS format and converted to tabular CSV format. The final dataset comprised digitisations of 16 landmarks on each of 339 male *L. botrana* forewings, grouped by adult flight season.

**Figure 1.**
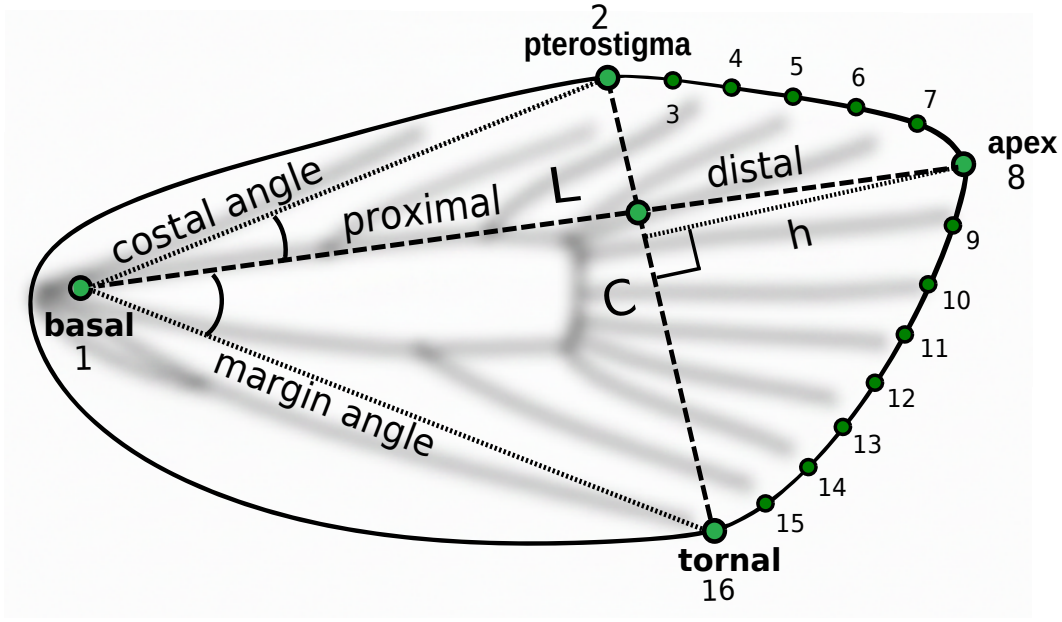
Digitisation scheme and basic metrics of the forewing of the European grapevine moth *Lobesia botrana*. Landmarks (green dots): 1—wing base (radius); 2—apex of pterostigma; 3–15—semilandmarks along the distal margin; 16—tornus A(1+2). *L*—longitudinal axis (length); *C*—chord (width); *h*—height from apex to chord.

### Data analysis

To characterise wing-planform variation, we developed MorphoGet, an open-source program [55] implemented as a Python class that adds flight-related morphological metrics (hereafter metrics) to a table of landmark coordinates. An executable Jupyter Notebook that reproduces the analyses is available in the repository [56], along with the data and the source code. A PDF of the executed notebook, together with Figures S1 and S2, is included in the Supplementary Materials. The calculation procedures are described below.

Generalised Procrustes Analysis (GPA) was used to align landmarks, removing translation, rotation, and scale; the resulting Procrustes coordinates were projected into tangent space [45, 57, 58]. Elliptic Fourier Analysis (EFA) was performed using PyEFD [59, 60]. We selected 15 EFA harmonics following the principle described in [47], balancing contour-description accuracy against the inclusion of high-frequency noise. This number provided an adequate visual approximation of wing shape without visible artifacts; using a different number of harmonics did not materially affect the results. Principal component analysis (PCA) and component deflation [61] were performed using scikit-learn [62, 63].

For GPA, the first component, interpreted as digitisation error (random error) [49], was removed by deflation. For EFA, coefficients were computed from raw, unaligned landmarks, and the first principal component, interpreted as the size component, was removed. PCA was then repeated for both datasets. Because principal-component signs are arbitrary, their orientations were fixed consistently based on the biplots and regressions onto landmarks.

Variation was visualised by regressing the metrics onto landmarks [44, 64]. For clarity, adjacent landmarks were connected with line segments around the wing outline. For visu-alisations of the metrics and GPA principal components, we used Matplotlib’s [65] smooth parameter; the degree of smoothing was selected for visual clarity. All parameters were visualised using GPA-aligned landmarks.

Pearson correlation coefficients were calculated using pandas [66] and used to assess the correspondence between the metrics and principal components. Levene’s test was used to assess the homogeneity of variance. Mean differences were tested using Student’s *t*-test when the assumption of equal variances was not rejected and Welch’s *t*-test otherwise. Statistical calculations were performed using SciPy [67].

### Computation of flight metrics

To simplify digitisation and computation, three metrics—aspect ratio (AR), a proxy for the second moment of wing area (MA), and sweep angle (SA)—were computed from four selected landmarks (landmarks 1, 2, 8, and 16). The wing base (landmark 1) and the most distal point of the bounding curve, the wing apex (landmark 8), define the longitudinal axis (*L*; Figure 1). The pterostigma apex (landmark 2) and the point where the anal vein meets the wing margin (landmark 16) define the chord (*C*), an operational measure of width that essentially corresponds to the broadest portion of the forewing.

Aspect ratio was calculated as wing length divided by width (AR = *L/C*) [26, 28]. The position of chord *C* along the longitudinal axis *L* serves as a simple marker of spanwise area distribution and is treated here as a proxy for the second moment of wing area (MA) [9]. MA was derived as the second principal component (PC2) of a PCA on the proximal and distal segments of *L*, which are separated by chord *C*. The first component represents overall axis length, whereas the second captures the inverse relationship between the segment lengths—that is, the displacement of the chord along the axis.

The sweep-angle metric is based on the inclination of the longitudinal axis *L* (from the wing base, landmark 1, to the apex, landmark 8) relative to the chord endpoints (landmarks 2 and 16). Like MA, SA was derived as PC2 of a PCA, here performed on the costal and marginal angles. These angles describe the relative displacement of the chord endpoints from the longitudinal axis. The first component represents chord size and its position relative to the wing base, corresponding to the size of the proximal wing region. For biplot visualisation, MA and SA measurements were standardised to a mean of 0 and a variance of 1 before PCA (*z*-transformation) [68, 69].

Apex taper (Taper) was calculated from the contour landmarks (landmarks 2–16). This adapted computational-morphology index [27] measures how the distal wing contour differs from a reference triangle with base *C* and height *h*, where *h* is the distance from the wing apex to the chord. The area of the distal wing segment (*S*_taper_) was normalised by the area of the reference triangle, *S*_max_ = 0.5 *· C · h*, to obtain *K*_raw_ = *S*_taper_*/S*_max_. For a fixed base andheight, the segment area ranges from that of a triangle (*K*_raw_ = 1) to that of a semi-ellipse (*K*_raw_ = *π/*2). We linearly transformed this ratio for ease of interpretation:

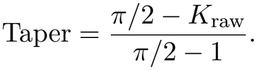

Thus, Taper = 1 corresponds to an ideal sharp triangle (maximum taper), whereas Taper = 0 corresponds to a semi-ellipse (minimum taper, or the bluntest shape).

## Results

The first two principal components derived from the PCA of the distal and proximal segments have a clear geometric interpretation. PC1 reflects overall wing length (L) — the combined length of the distal and proximal segments, as indicated by the loading-vector directions (Figure S1, left). PC2 captures the inverse relationship between these segments, reflecting a shift in the position of chord C along the longitudinal axis. The costal and marginal angles show a similar inverse relationship (Figure S1, right). The metrics computed from four landmarks thus represent interpretable geometric features: MA tracks the position of the chord along the longitudinal axis, and SA tracks the inclination of the longitudinal axis relative to the chord.

Landmark 2 was placed by visual judgment at the top of the pterostigma (landmark type 2), which introduces positional shift error [49]. A small displacement of this anchor point shifts the entire landmark chain (landmarks 2–16), which appears in visualisations as a “rotation” of the wing outline (Figure S2 (left)). After this error component was removed by deflation, PCA was repeated and five principal components were extracted. In the EFA of raw, unaligned landmarks, the first principal component corresponds to overall size (Figure S2 (right)). After this component was removed, the repeated PCA also yielded five components. Figure 2 shows the principal components from GPA and EFA, together with the flight-related morphometric variables and their contributions to shape variation.

**Figure 2.**
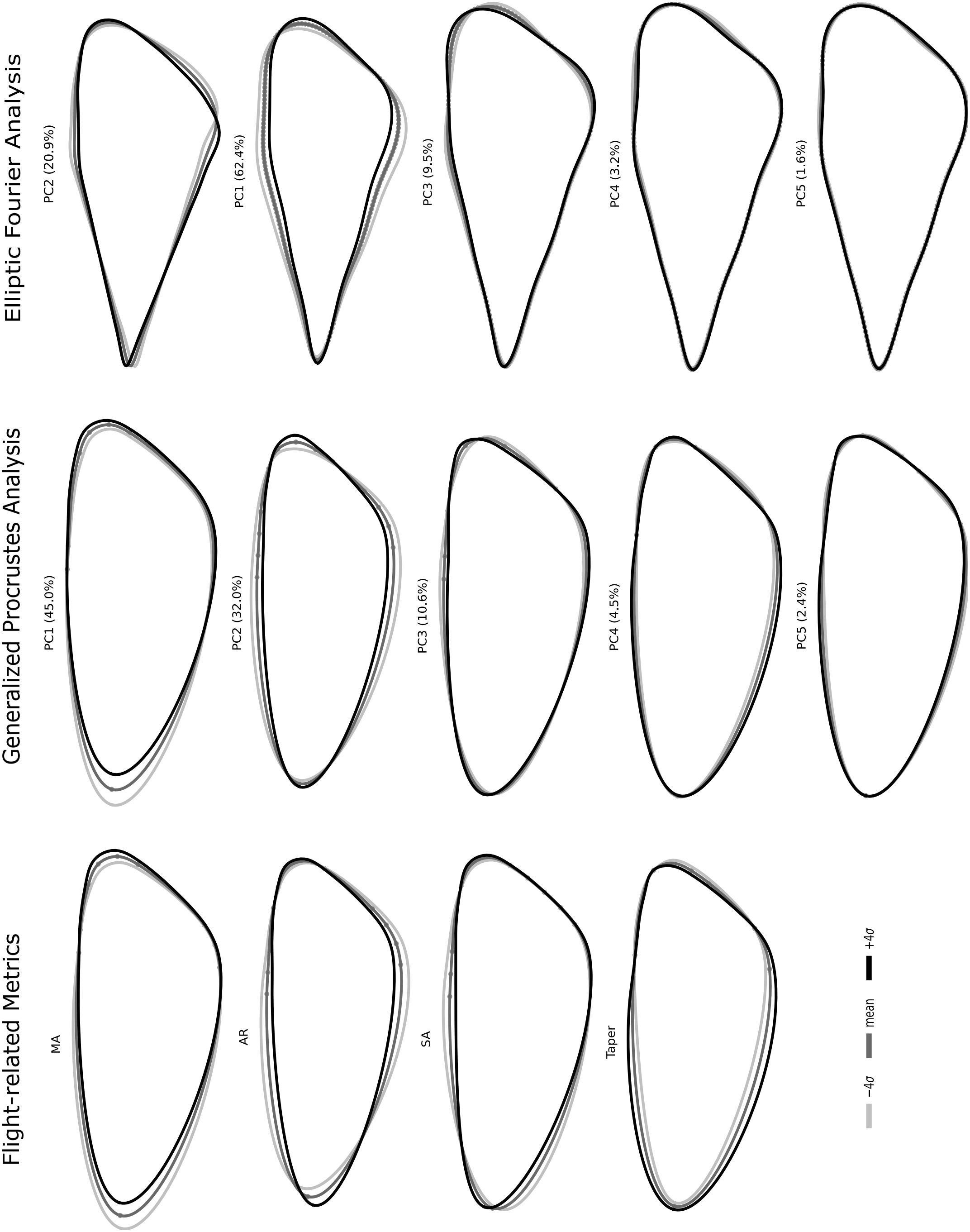
Visualisation of flight-related morphological metrics and GPA / EFA principal components. Lines show the mean shape and deviations from it (*±*4*σ*). Grey line: *™*4*σ*; dark grey line: mean; black line: +4*σ*.

Table 1 reports the correlations between the metrics and the principal components of GPA and EFA. For both methods, each of the first four components showed its strongest absolute correlation with a different metric, but the order differed between methods: in GPA, PC1 with MA and PC2 with AR; in EFA, PC1 with AR and PC2 with MA; and in both, PC3 with SA and PC4 with Taper. The three metrics calculated from four homologous landmarks yielded components that accounted for 87.8% of the analysed variation in GPA and 92.5% in EFA; PC4, associated with Taper, contributed a further 4.5% and 3.2%, respectively, and captured variance in a localised region of the wing. Regression visualisations of the fifth GPA and EFA components suggest a similar pattern, resembling sweep-angle variation but confined to the distal wing (2.4% and 1.6% of variation, respectively). The correlations for PC5.were not tested.

**Table 1.** Correlations of flight-related morphological metrics with GPA and EFA principal components (*n* = 339). Cell colour intensity is proportional to |*r*|. The strongest correlation for each component (highlighted in bold) identifies the metric to which that component corresponds. Component and metrics contributions to variation are shown in Figure 2.

| **Component** | **MA** | **AR** | **SA** | **Taper** |
|---|---|---|---|---|
| GPA PC1 | **-0.933**<sup>\*\*\*</sup> | −0.419<sup>***</sup> | −0.492<sup>***</sup> | −0.308<sup>***</sup> |
| GPA PC2 | −0.287<sup>***</sup> | **0.775**<sup>\*\*\*</sup> | −0.125<sup>*</sup> | −0.281<sup>***</sup> |
| GPA PC3 | −0.137<sup>*</sup> | 0.095 | **-0.787**<sup>\*\*\*</sup> | 0.369<sup>***</sup> |
| GPA PC4 | 0.018 | 0.255<sup>***</sup> | −0.123<sup>*</sup> | **-0.707**<sup>\*\*\*</sup> |
| **Component** | **AR** | **MA** | **SA** | **Taper** |
| EFA PC1 | **-0.743**<sup>\*\*\*</sup> | −0.402<sup>***</sup> | −0.179<sup>**</sup> | 0.147<sup>**</sup> |
| EFA PC2 | −0.431<sup>***</sup> | **0.582**<sup>\*\*\*</sup> | 0.464<sup>***</sup> | 0.035 |
| EFA PC3 | 0.015 | −0.045 | **0.688**<sup>\*\*\*</sup> | −0.192<sup>***</sup> |
| EFA PC4 | −0.247<sup>***</sup> | −0.131<sup>*</sup> | −0.118<sup>*</sup> | **0.612**<sup>\*\*\*</sup> |

*Note.* <sup>\*</sup> *p* < 0.05; <sup>\*\*</sup> *p* < 0.01; <sup>\*\*\*</sup> *p* < 0.001. MA—proxy for the second moment of wing area; AR—aspect ratio; SA—sweep angle; Taper—apex taper. Cell colour intensity is proportional to \|*r*\| (scale: white = 0, saturated red = 1). Bold values mark the strongest correlation for each principal component.

Levene’s test indicated unequal variances between generations for Taper but not for MA, AR, or SA (Table 2). Accordingly, means were compared using Welch’s *t*-test for Taper and Student’s *t*-test for the other metrics. The spring and summer generations differed significantly in MA, AR, and SA (all p<0.001), but not in Taper (p=0.081; Table 2). The spring sample was half the size of the summer sample (n=112 vs. n=227) and had a larger standard deviation for Taper (0.0542 vs. 0.0410), indicating greater dispersion (Figure 3).

**Table 2.**
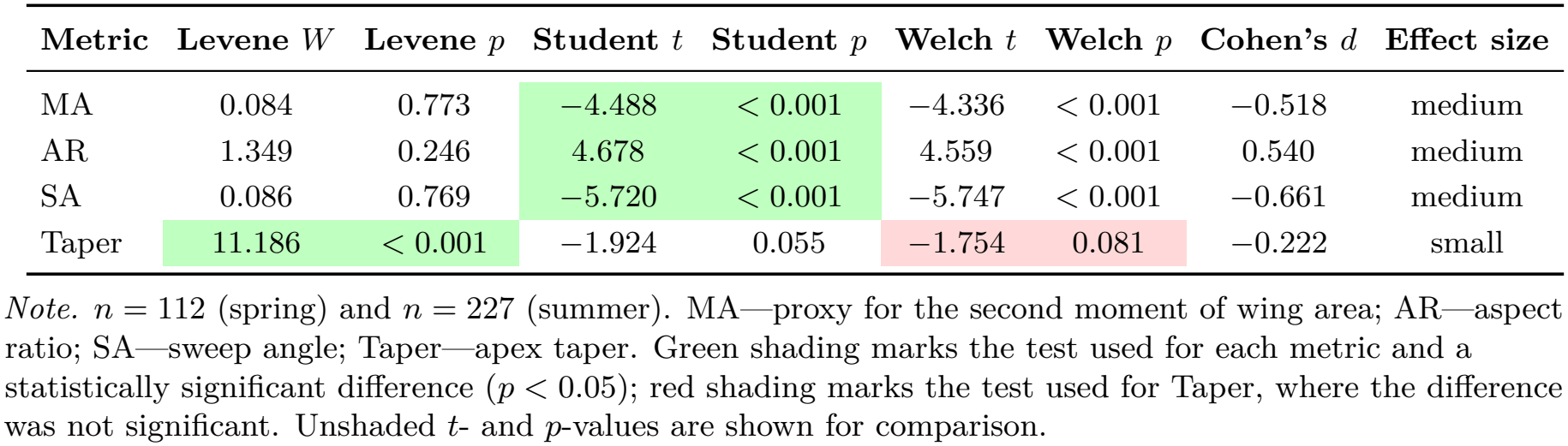
Comparison of flight-related metrics between seasonal generations (spring vs. summer) of *Lobesia botrana*. Levene’s test assessed homogeneity of variance. For MA, AR, and SA, equal variances were not rejected, and Student’s *t*-test was used (Welch’s *t*-test shown for comparison). For Taper, variances were unequal, and Welch’s *t*-test was used (Student’s *t*-test shown for comparison). Effect sizes are given as Cohen’s *d*.

| Metric | Levene *W* | Levene *p* | Student *t* | Student *p* | Welch *t* | Welch *p* | Cohen's *d* | Effect size |
|---|---|---|---|---|---|---|---|---|
| MA | 0.084 | 0.773 | −4.488 | < 0.001 | −4.336 | < 0.001 | −0.518 | medium |
| AR | 1.349 | 0.246 | 4.678 | < 0.001 | 4.559 | < 0.001 | 0.540 | medium |
| SA | 0.086 | 0.769 | −5.720 | < 0.001 | −5.747 | < 0.001 | −0.661 | medium |
| Taper | 11.186 | < 0.001 | −1.924 | 0.055 | −1.754 | 0.081 | −0.222 | small |

*Note.* *n* = 112 (spring) and *n* = 227 (summer). MA—proxy for the second moment of wing area; AR—aspect ratio; SA—sweep angle; Taper—apex taper. Green shading marks the test used for each metric and a statistically significant difference (*p* < 0.05); red shading marks the test used for Taper, where the difference was not significant. Unshaded *t*- and *p*-values are shown for comparison.

**Figure 3.**
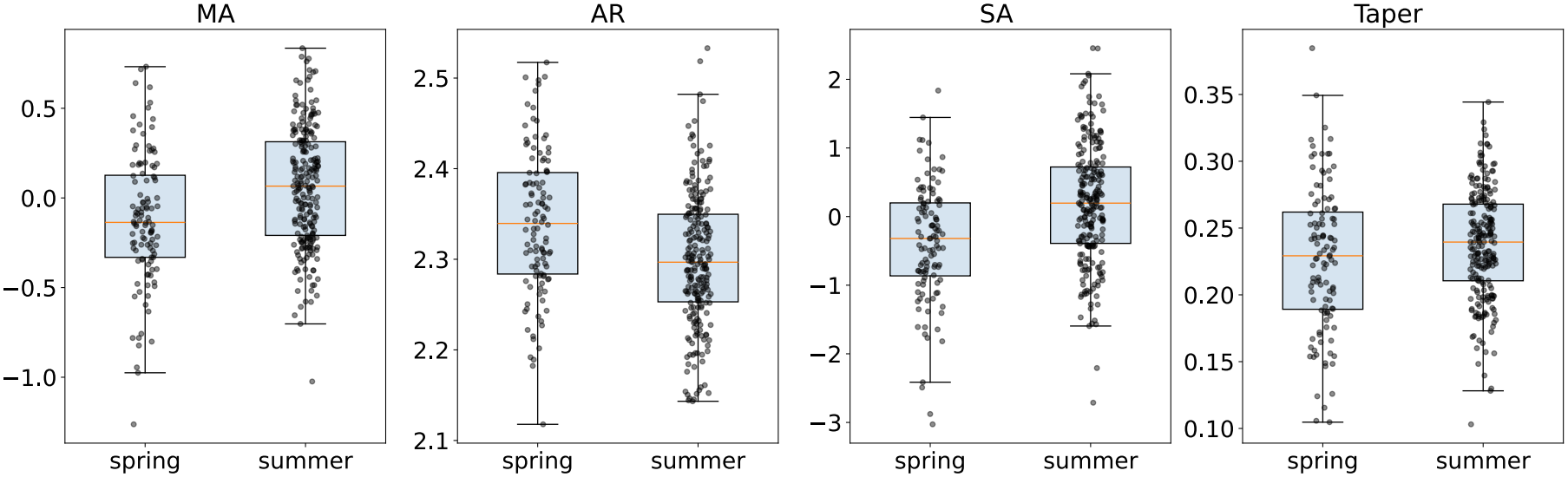
Boxplots of flight-related wing metrics for the spring and summer generations. MA—proxy for the second moment of wing area; AR—aspect ratio; SA—sweep angle; Taper—apex taper.

## Discussion

GPA removes scale during alignment, and the first size-related EFA component was removed by deflation (estimating the size component and its contribution was not an objective of this study). The metrics and principal components were derived from the same landmarks, so some correlation — particularly in the case of GPA — is inherent in the analytical design. However, at this stage, we conclude only that the selected metrics adequately capture the identified patterns of variation and correspond to the underlying variation. The first four metric-associated components accounted for 92.3% (GPA) and 95.7% (EFA) of the analysed shape variation. If PC5 is also considered, only 5.3% of the GPA variance and 2.7% of the EFA variance remain unexplained. These proportions are small relative to the variance typically discarded in standard PCA after GPA [70, 71] and should largely reflect the cumulative effects of digitisation/computational errors (including nonlinear relationships) and developmental instability. There is therefore little or no remaining variation, whether phenotypic plasticity or genetically related variability.

The principal components obtained by GPA and EFA showed moderate to strong correlations with flight-related metrics (Table 1), and regressions of metrics and components onto landmarks revealed similar patterns of wing-planeform variation (Figure 2) [64]. The two complementary methods of shape description yielded concordant results and similar visualisations, supporting the robustness of the identified patterns. Seasonal-generation differences are environmental (non-genetic) in origin — a case of seasonal polyphenism reflected in these metrics. Similar seasonal differences in AR and MA have been reported in Lepidoptera, although AR and apex taper can be difficult to distinguish [72–74]. These differences could be associated with different flight demands, and this distinction is important to assess the ecological and adaptive status of populations [12, 31, 32, 75–77]. Together, these findings indicate that a small set of flight-related morphological metrics captures major axes of complex wing-planform plasticity with both methods. Specifically, GPA and EFA revealed five patterns (the first five principal components) of wing-planform variation in male *L. botrana*, four of which were biologically meaningful in correlation analysis (AR, MA, SA, and Taper). Three of these — AR, MA, and SA — capture whole-wing shape and are calculated from four homologous landmarks on the wing surface. However, it seems that similar key points could be found for the other patterns as well.

The correspondence between the first principal component and aspect ratio has previously been reported in dragonflies [78]. Re-examination of published disparity visualisations also suggests that aspect ratio and the second moment of wing area can be discerned in other studies [6, 79–81]. A change in area distribution is likewise apparent in published plots of beetle miniaturisation within a single genus, although the authors described it in terms of aspect ratio [82]. Studies of wing-shape variation across major clades of Hymenoptera and Diptera have also reported that the first two principal components correspond to aspect ratio and the second moment of wing area [83, 84]. Together, these studies suggest that such metrics may capture important axes of wing-planform variation at both the intraspecific and interspecific levels. Accounting for this structure may reveal otherwise obscured differences and help identify distinct functional strategies across groups.

In one of his last works, Fred L. Bookstein — one of the founders of geometric morphometric methods — provided a skeptical review of the applicability of landmark-based methods, citing a number of limitations [48]. This critique, far from undermining the value of these methods, reflects the ongoing development of the field and the rigor of its methodological debate. With these limitations in mind, and as our results show, biologically meaningful hypotheses can nevertheless be successfully inferred from empirical geometric patterns. Moreover, studying form as a whole offers distinct advantages: it captures both the integrated whole and its constituent parts — their relationships, contributions, and relative shares, as well as the residual variation. The selected metrics obviate the need to explore additional parameters in *Lobesia botrana* male forewing shape, except for localised variation in the proximal wing region (which was not digitised). It also clearly separates apex taper (Taper) from aspect ratio (AR). Ultimately, correlation analysis across different morphometric methods and its biological significance allowed us to establish morphological metrics as biologically meaningful features that are informative in their own right. These findings offer a solid foundation for further morphological and functional investigation of these characters and their interrelationships — as well as for a broad range of ecological and evolutionary research — including how these parameters relate to insect behaviour and development.

In practice, the approach is remarkably simple: four homologous landmarks spanning the wing planform proved sufficient for calculating metrics without labour-intensive procedures such as scale removal or vein-junction digitisation, and without complex multivariate analyses. This streamlined approach may facilitate the wider use of morphological characters as markers of wing shape. However, the relationship between metrics and wing shape should be assessed separately for each insect group. These features may differ among insect taxa due to structural and functional factors — for example, the organisation of flapping flight (bipartite vs. unipartite), wing sclerotisation, and functional specialisation (e.g., the forewings of some crickets, the hemelytra of true bugs, wing-folding mechanisms). Such differences are no less informative — they can provide a deeper understanding of how structure underlies function and shapes trade-offs between functional demands. The metrics can be derived for other taxa using the open-source MorphoGet code: it can be applied directly or, where wing-shape variation differs or is more complex, serve as a foundation for formal analyses.

## Supporting information

Electronic Supplementary Materials. 2.Jupyter Notebook

Electronic Supplementary Materials. 1.Figures

## Data and code availability

### Data

The raw digitised landmark dataset (CSV) and the executable Jupyter Notebook that reproduces all calculations are available in the open repository Zenodo [56]. A PDF version of the executed notebook and a separate PDF with Figures S1 and S2 are provided in the Supplementary Materials.

### Code

The MorphoGet program (v.0.1.0) is available in the repository [55]; the complete analysis pipeline — GPA, EFA, deflation, computation of flight-related morphological metrics, statistics, and visualisation — is implemented as an executable Jupyter Notebook [56].

## Author contributions

**Oleg V. Orlov:** Conceptualisation; Methodology; Investigation; Data Curation; Formal Analysis; Software; Visualisation; Writing – original draft. **Evgeniya G. Yurchenko:** Supervision; Writing – review & editing.

