## Supplementary material for "A small set of morphological metrics describes wing-planform plasticity: a correlational analysis in a population of the European grapevine moth *Lobesia botrana* (Lepidoptera: Tortricidae)": Electronic Supplementary Materials. 2.Jupyter Notebook

November 24, 2026

### 0.0.1 Program start and data loading

[4]:
```
from MorphoGet import MorphoGet
get = MorphoGet.from_csv("Lobesia botrana.csv")
```

```
Data loaded from 'Lobesia botrana.csv'. Rows: 339
```

### 0.0.2 Assigning landmarks

[5]:
```
get.set_landmarks()
get.plot_consensus(figsize=(6, 4))
get.set_fa_params(Apex=8, Basal=1, Costa=2, Tornal=16)
get.plot_schematic(figsize=(8, 5))
```

```
Selected 16 points.
```

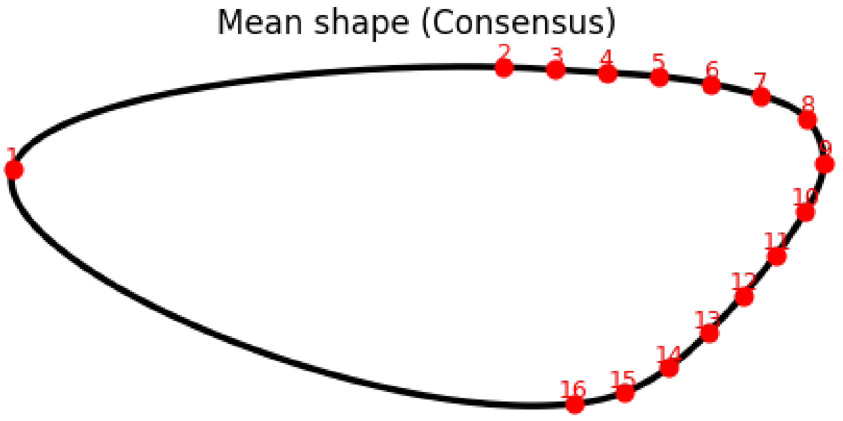


```
'Apex' -> point 8
'Basal' -> point 1
'Costa' -> point 2
'Tornal' -> point 16
```

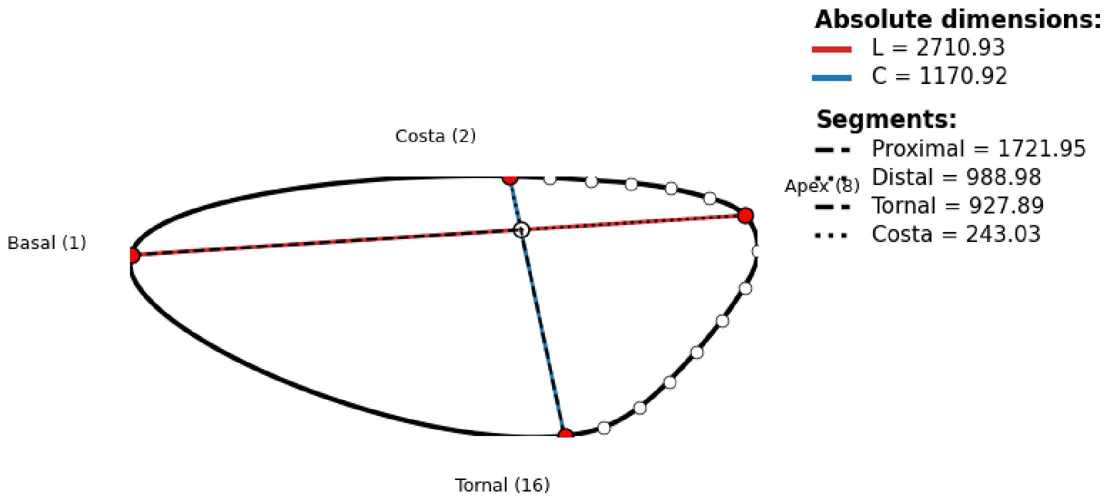


### 0.0.3 alculate Flight-related Metrics

[3]:
```
get.calculate_fa_metrics(ma_l_comp=1, sa_a_comp=1, taper_l_comp=1)
get.plot_biplot('ma_biplot', 'sa_a_biplot')
```

```
FA calculated (MA_l=1, SA_l=1, SA_a=1, Taper_l=1): MA_l, MA_s, AR_l, AR_s,
SA_l, SA_a, Taper_l, Taper_K
```

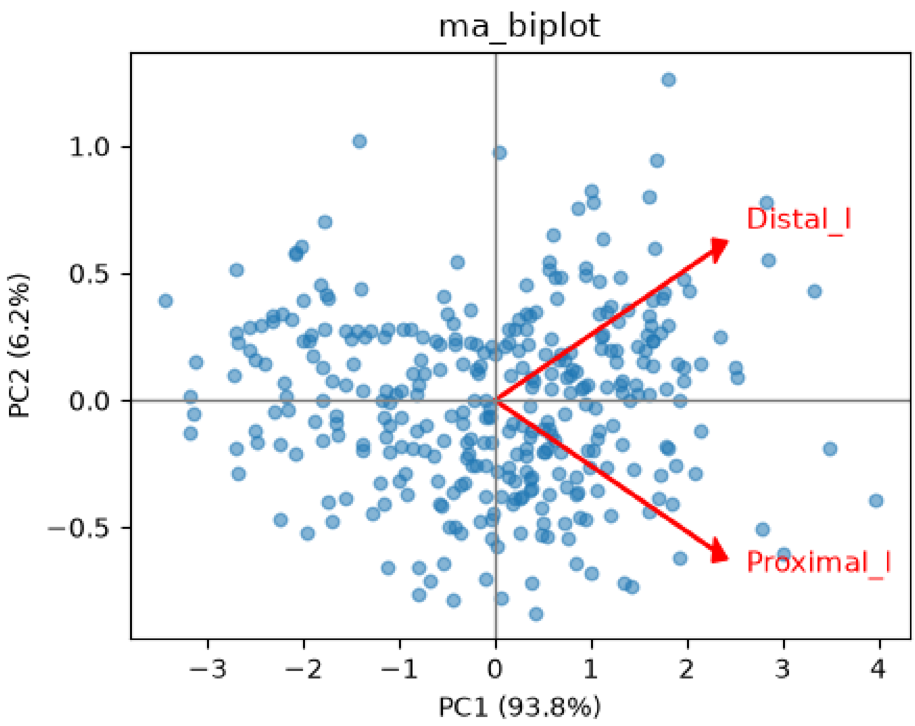

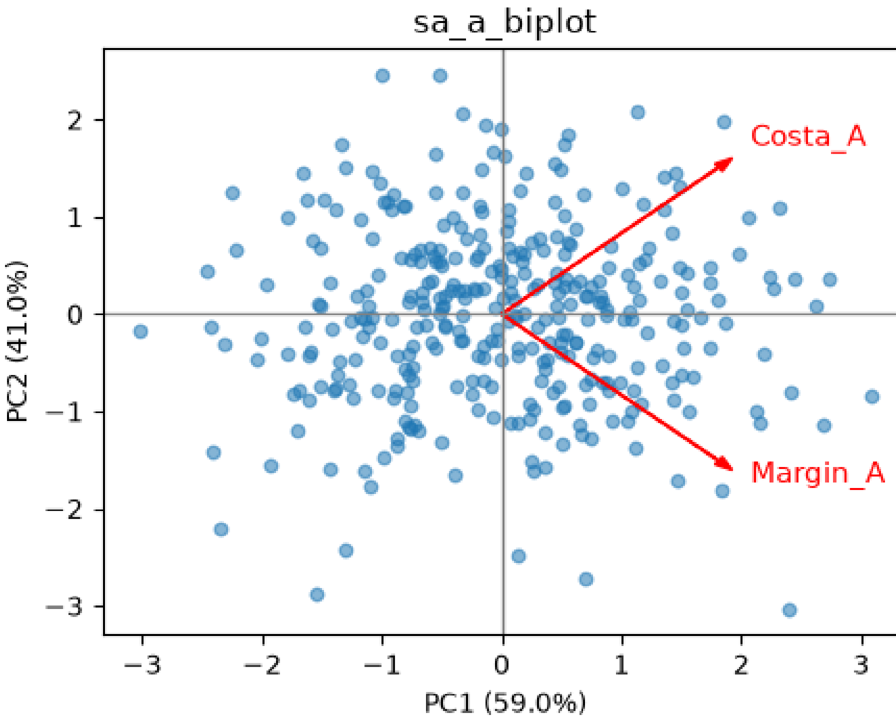


[4]:
```
get.run_gpa()
get.plot_regressions(fa_metrics=['MA_l', 'AR_l', 'SA_a', 'Taper_K'], n_cols=2,␣
↪z_ext=4.0, smooth=True, invert_sigma=['SA_a'])
```

```
GPA completed.
```

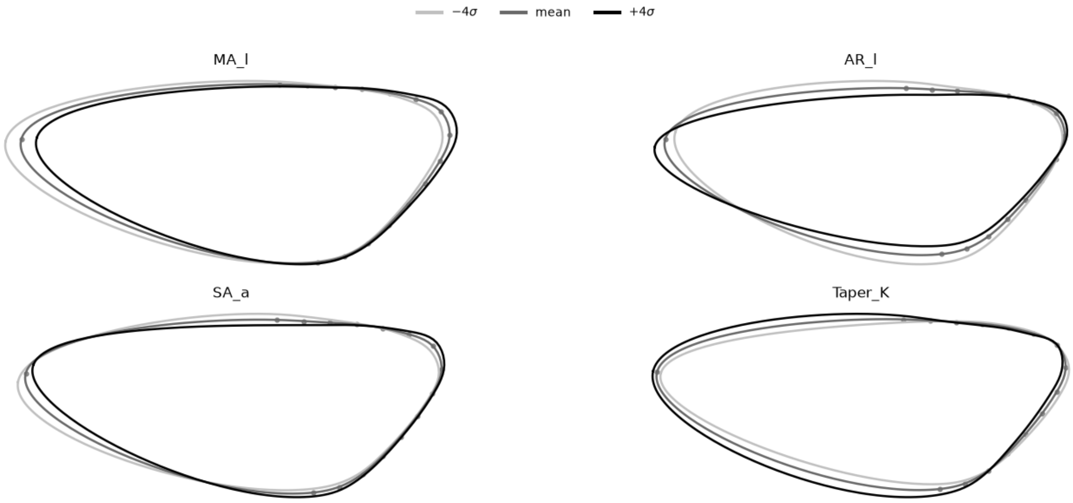

### 0.0.4 Generalized Procrustes Analysis (GPA)

[5]:
```
get.plot_pca(pc_indices=[0], n_cols=2, smooth=True, z_ext=4.0)
```

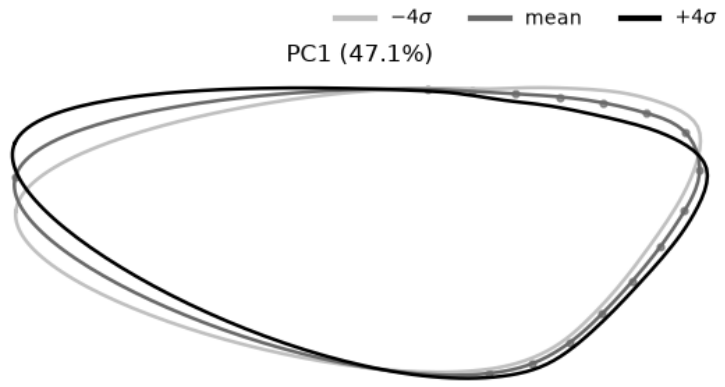


[6]:
```
get.neutralize_pcs(components_to_remove=[0])
get.plot_pca(pc_indices=[0, 1, 2, 3, 4], n_cols=2, z_ext=4.0, smooth=True,
 invert_sigma=[3])
```

```
Removed PC1 (47.1%)
```

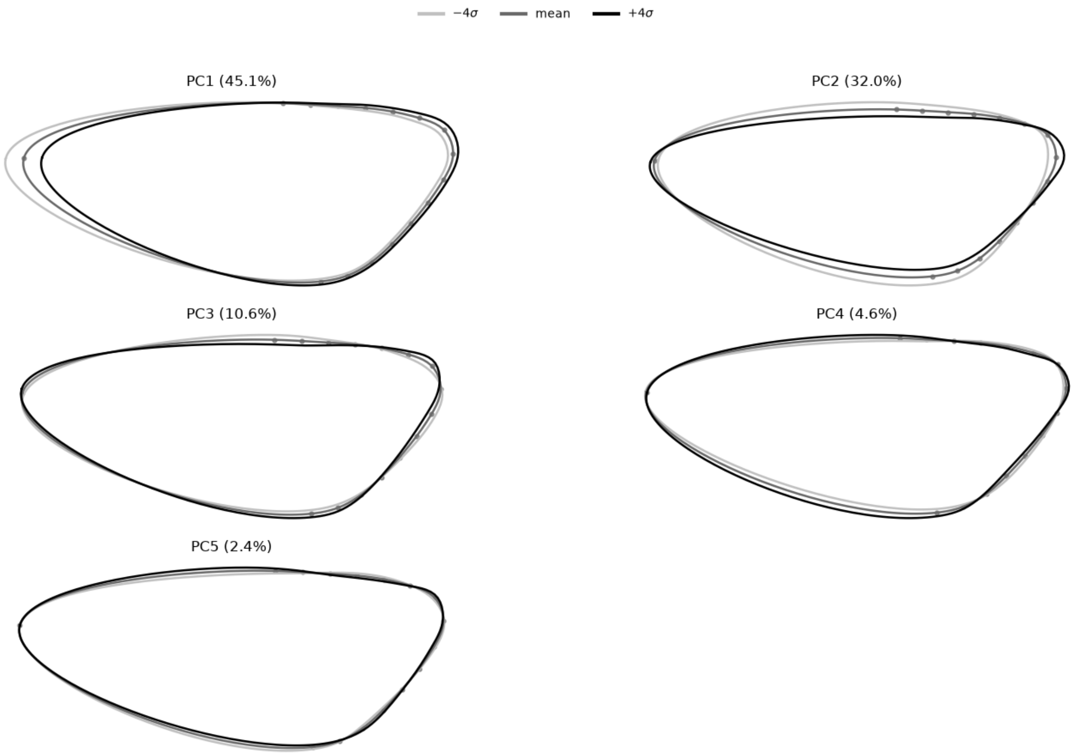

[7]:
```
get.show_correlations(['MA_l', 'AR_l', 'SA_a', 'Taper_K'])
```

```
<pandas.io.formats.style.Styler at 0x1d753a12b90>
```

### 0.0.5 Elliptic Fourier Analysis (EFA)

[8]:
```
get.calculate_efa(order=15, normalize=True, use_neutralized=False)
get.run_efa_pca(n_components=1)
get.plot_efa_pca(pc_indices=[0], n_cols=2, z_ext=0.5, flip_x=True, flip_y=True)
```

```
 EFA calculated: 339 contours, order=15, normalize=True
 PCA on EFA completed. Explained variance ratio: [0.997]
```

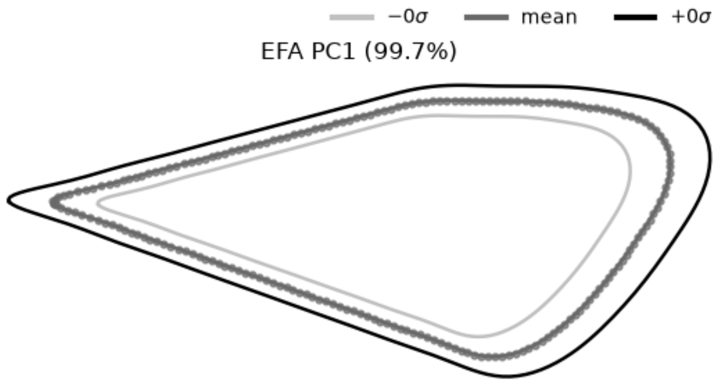


[9]:
```
get.calculate_efa(order=15, normalize=True, use_neutralized=False)
get.neutralize_pcs(components_to_remove=[0])
get.run_efa_pca(n_components=6)
get.plot_efa_pca(pc_indices=[0, 1, 2, 3, 4], n_cols=2, z_ext=4.0,␣
↪invert_sigma=[3], flip_x=True, flip_y=True)
```

```
 EFA calculated: 339 contours, order=15, normalize=True
Removed EFA PC1 (99.7%)
 PCA on EFA completed. Explained variance ratio: [0.624 0.205 0.095 0.033 0.017
0.009]
```

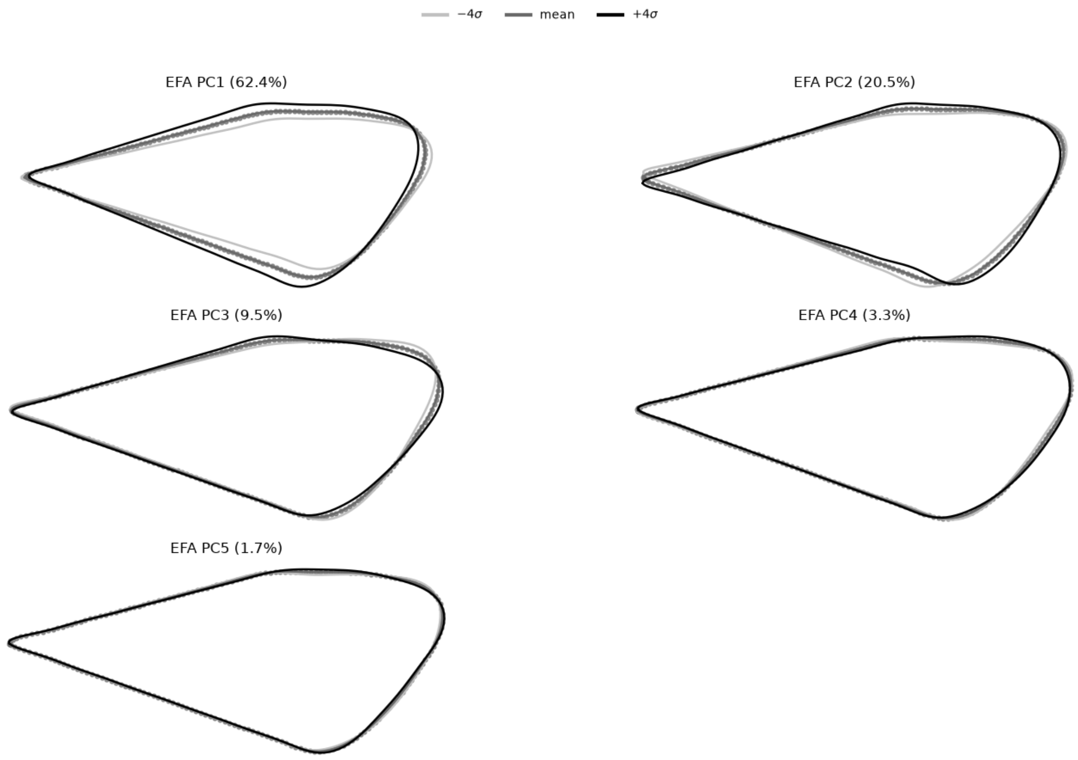


[10]:
```
get.show_efa_correlations(['AR_l', 'MA_l', 'SA_a', 'Taper_K'], n_efa_pcs=4)
```

```
<pandas.io.formats.style.Styler at 0x1d7533f7310>
```

### 0.0.6 Group comparison

[11]:
```
import numpy as np
import pandas as pd
import matplotlib.pyplot as plt
from scipy import stats

def cohens_d(x, y):
    nx, ny = len(x), len(y)
    pooled_sd = np.sqrt(((nx - 1) * x.var(ddof=1) + (ny - 1) * y.var(ddof=1)) /␣
 ↪(nx + ny - 2))
    return np.nan if pooled_sd == 0 else (x.mean() - y.mean()) / pooled_sd

def interpret_d(d):
    ad = abs(d)
    if np.isnan(ad):
        return "NA"
    if ad < 0.2:
        return "negligible"
```

```python
    if ad < 0.5:
        return "small"
    if ad < 0.8:
        return "medium"
    return "large"

def compare_two_groups(mg, group_col, metrics, groups, alpha=0.05,
                       annotate=True, figsize=None, save_path=None, dpi=300):
    fmt_p = lambda p: f"{p:.4f}" if p >= 0.001 else "<0.001"

    df = mg.current_df
    g1 = df[df[group_col] == groups[0]]
    g2 = df[df[group_col] == groups[1]]

    rows = []
    for m in metrics:
        x = g1[m].dropna()
        y = g2[m].dropna()

        lev = stats.levene(x, y)
        stud = stats.ttest_ind(x, y, equal_var=True)
        welch = stats.ttest_ind(x, y, equal_var=False)
        d = cohens_d(x, y)

        rows.append({
            'metric': m,
            'n': f"{len(x)}/{len(y)}",
            f'mean±SD ({groups[0]})': f"{x.mean():.3f}±{x.std():.3f}",
            f'mean±SD ({groups[1]})': f"{y.mean():.3f}±{y.std():.3f}",
            'Levene W': f"{lev.statistic:.3f}",
            'Levene p': fmt_p(lev.pvalue),
            'Student t': f"{stud.statistic:.3f}",
            'Student p': fmt_p(stud.pvalue),
            'Welch t': f"{welch.statistic:.3f}",
            'Welch p': fmt_p(welch.pvalue),
            "Cohen's d": f"{d:.3f}" if not np.isnan(d) else "NA",
            'Effect size': interpret_d(d),
        })

    table = pd.DataFrame(rows)

    print(f"Two-group comparison by '{group_col}': {groups[0]} vs {groups[1]}␣
↪(alpha={alpha})")
    print(table[['metric', 'Levene W', 'Levene p',
                 'Student t', 'Student p',
                 'Welch t', 'Welch p',
                 "Cohen's d", 'Effect size']].to_string(index=False))
```

```
    n = len(table)
    TITLE_SIZE, TICK_SIZE = 22, 20
    fig, axes = plt.subplots(1, n, figsize=figsize or (4.8 * n, 6.0),␣
↪squeeze=False)

    for ax, (_, row) in zip(axes[0], table.iterrows()):
        m = row['metric']
        x = g1[m].dropna()
        y = g2[m].dropna()

        bp = ax.boxplot([x, y], showmeans=False, showfliers=False,
                        patch_artist=True, widths=0.5)
        for box in bp['boxes']:
            box.set_facecolor('#d6e4f0')

        for i, data in enumerate([x, y], start=1):
            jitter = np.random.uniform(-0.08, 0.08, size=len(data))
            ax.scatter(np.full(len(data), i) + jitter, data,
                       color='black', alpha=0.45, s=18, zorder=3)

        ax.set_xticks([1, 2])
        ax.set_xticklabels(groups, fontsize=TICK_SIZE)
        ax.tick_params(axis='y', labelsize=TICK_SIZE)
        ax.set_title(m, fontsize=TITLE_SIZE)

    plt.tight_layout()

    if save_path:
        if not save_path.endswith('.svg'):
            save_path = save_path.rsplit('.', 1)[0] + '.svg'
        plt.savefig(save_path, bbox_inches='tight', format='svg', dpi=dpi)

    plt.show()
    return table
```

[12]:
```
table = compare_two_groups(
    mg=get,
    group_col='Generation',
    metrics=['MA_l', 'AR_l', 'SA_a', 'Taper_l'],
    groups=['spring', 'summer'],
    save_path='boxplots.png'
)
```

```
Two-group comparison by 'Generation': spring vs summer (alpha=0.05)
 metric Levene W Levene p Student t Student p Welch t Welch p Cohen's d Effect
size
   MA_l     0.084   0.7726     4.488    <0.001   4.336  <0.001     0.518
```

```
medium
   AR_l     1.349    0.2463     4.678     <0.001    4.559  <0.001      0.540
medium
   SA_a     0.086    0.7689    -5.720     <0.001   -5.747  <0.001     -0.661
medium
Taper_l    22.925    <0.001    -3.535     <0.001   -3.097  0.0023     -0.408
small
```

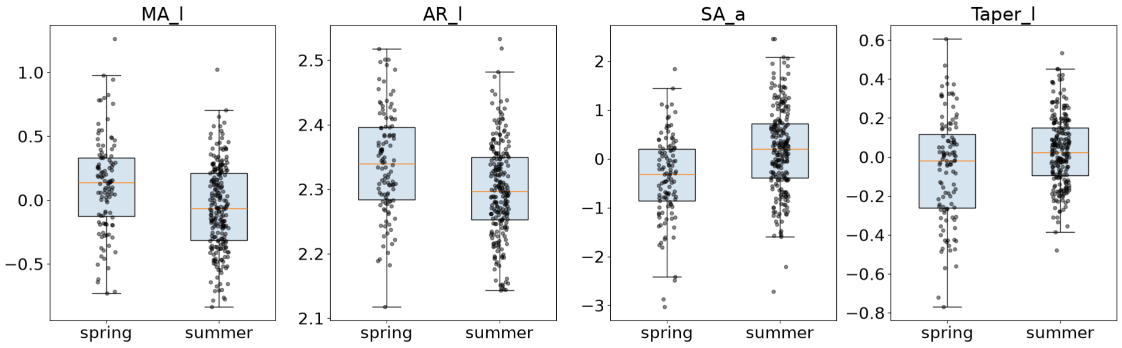


### 0.0.7 Grouped PCA biplots

**Ellipse_std=2.0 approximates the 95% region for bivariate normal data.**

[13]:
```python
from pathlib import Path

from matplotlib.patches import Ellipse
from sklearn.decomposition import PCA
from sklearn.preprocessing import StandardScaler


def _ellipse(ax, x, y, color, n_std):
    vals, vecs = np.linalg.eigh(np.cov(x, y))
    vals = np.sqrt(np.clip(vals, 0, None))
    ax.add_patch(Ellipse(
        xy=(x.mean(), y.mean()),
        width=vals[0] * n_std * 2,
        height=vals[1] * n_std * 2,
        angle=np.degrees(np.arctan2(*vecs[:, 0][::-1])),
        facecolor=color, alpha=0.15, edgecolor=color, lw=2,
    ))


def _arrows(ax, loadings, columns, scale):
    for (x, y), name in zip(loadings * scale, columns):
        ax.arrow(0, 0, x, y, color='red', width=0.01,
                 head_width=0.10, length_includes_head=True)
```

```
        ax.text(x * 1.08, y * 1.08, name, color='red', fontsize=11,␣
↪fontweight='bold')


def plot_grouped_biplots(mg, presets, group_col, groups=None, figsize=(6, 5),
                         ellipse_std=2.0, arrow_scale=2.5, save_path=None,␣
↪dpi=300):
    groups = groups or mg.current_df[group_col].dropna().unique().tolist()
    colors = {g: plt.get_cmap('tab10')(i % 10) for i, g in enumerate(groups)}

    for preset in presets:
        columns = mg._BIPLOT_PRESETS[preset]
        data = mg.lengths_df[columns].dropna()

        X = StandardScaler().fit_transform(data.values)
        pca = PCA(n_components=2, random_state=42).fit(X)
        scores = pca.transform(X)
        loadings = pca.components_.T * np.sqrt(pca.explained_variance_)
        evr = pca.explained_variance_ratio_ * 100

        pts = pd.DataFrame(scores, index=data.index, columns=['PC1', 'PC2'])
        pts[group_col] = mg.current_df.loc[data.index, group_col]
        pts = pts.dropna(subset=[group_col])

        fig, ax = plt.subplots(figsize=figsize)

        for g in groups:
            g_pts = pts[pts[group_col] == g]
            c = colors[g]
            x, y = g_pts['PC1'].values, g_pts['PC2'].values
            ax.scatter(x, y, s=35, alpha=0.7, label=str(g), color=c,
                       edgecolors='white', linewidth=0.5)
            if len(g_pts) >= 3:
                _ellipse(ax, x, y, c, ellipse_std)

        _arrows(ax, loadings, columns, arrow_scale)

        ax.axhline(0, color='gray', lw=0.5, ls='--')
        ax.axvline(0, color='gray', lw=0.5, ls='--')
        ax.set_xlabel(f'PC1 ({evr[0]:.1f}%)', fontsize=11)
        ax.set_ylabel(f'PC2 ({evr[1]:.1f}%)', fontsize=11)
        ax.set_title(preset, fontsize=13)
        ax.legend(title=group_col, fontsize=9)
        fig.tight_layout()

        if save_path:
            stem = Path(save_path).with_suffix('')
```

```
            ext = Path(save_path).suffix or '.png'
            fig.savefig(f"{stem}_{preset}{ext}", bbox_inches='tight', dpi=dpi)
        plt.show()
        plt.close(fig)
```

[14]:
```
plot_grouped_biplots(
    mg=get,
    presets=['ma_biplot', 'sa_a_biplot'],
    group_col='Generation',
    groups=['spring', 'summer'],
    save_path='biplots.svg',
)
```

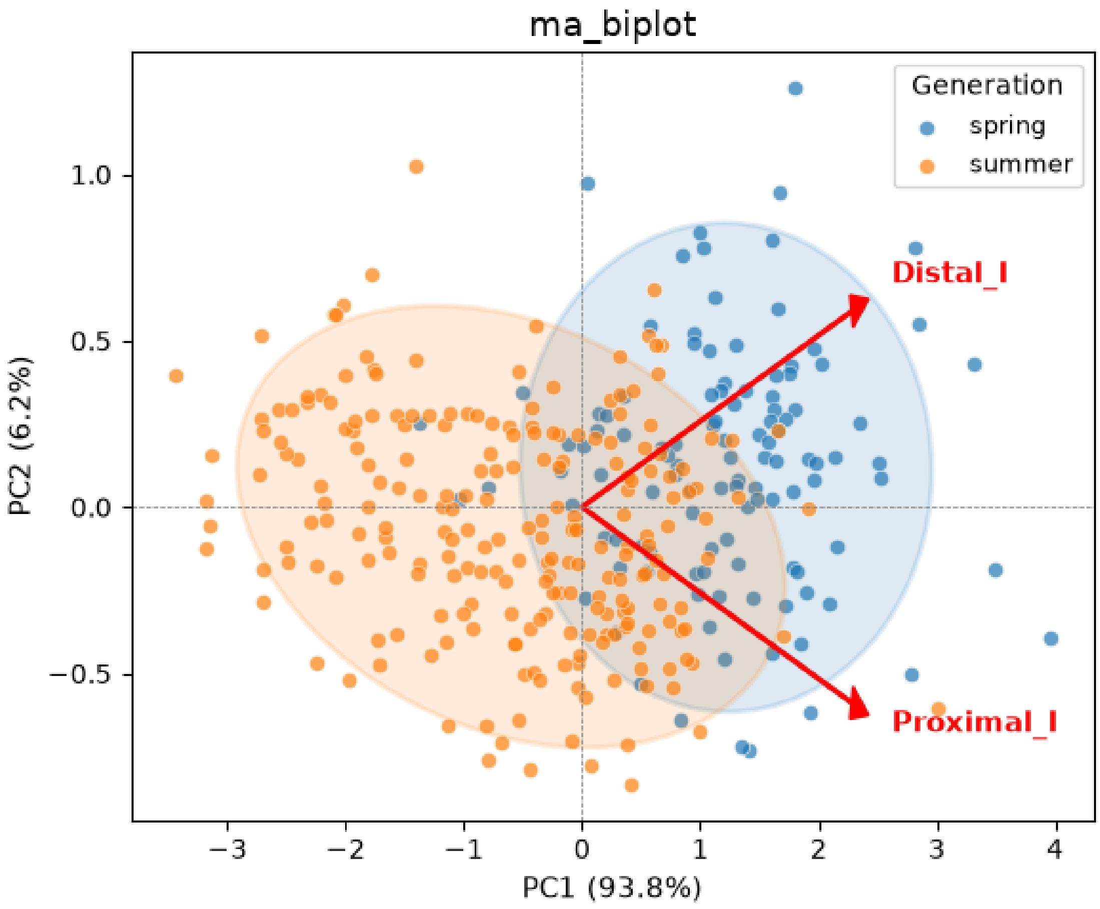

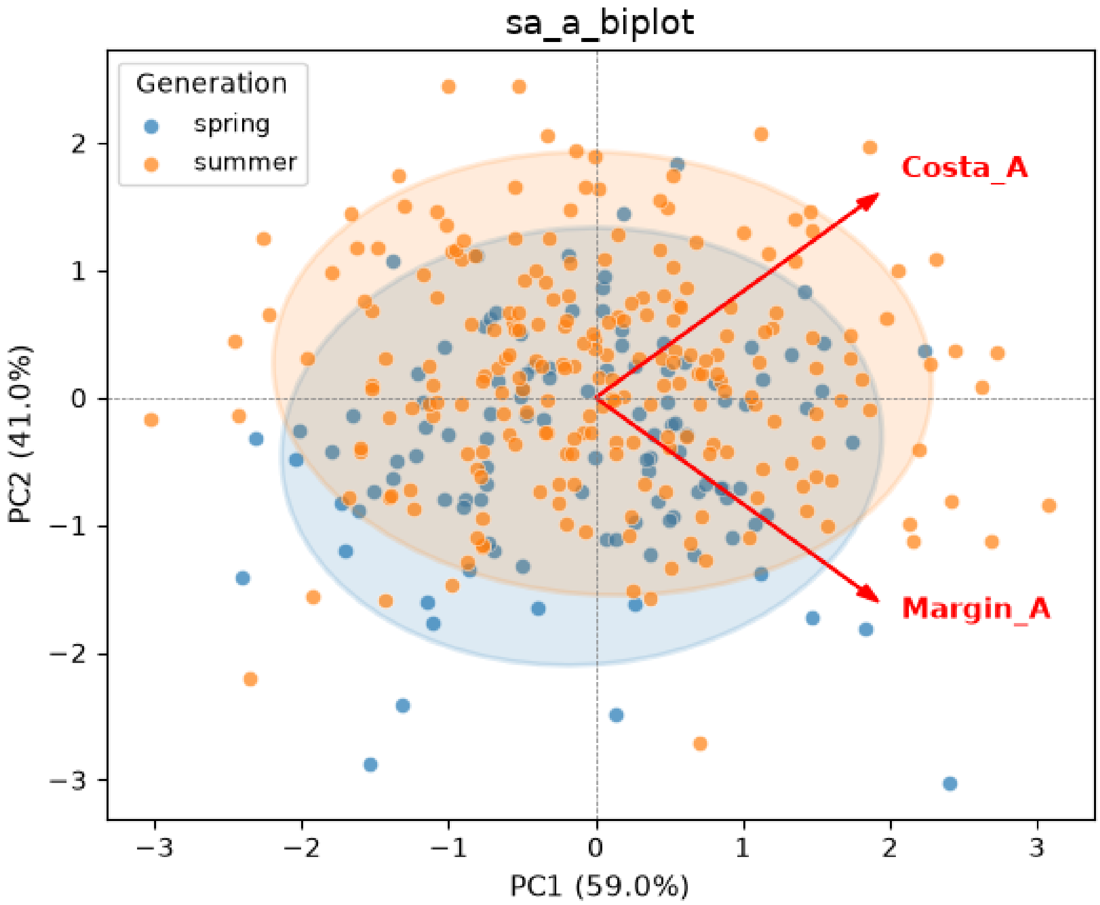
sa_a_biplot
Generation
spring
summer
Costa_A
Margin_A
PC1 (59.0%)
PC2 (41.0%)

[ ]:
