## Supplementary material for "A small set of morphological metrics describes wing-planform plasticity: a correlational analysis in a population of the European grapevine moth *Lobesia botrana* (Lepidoptera: Tortricidae)": Electronic Supplementary Materials. 1.Figures

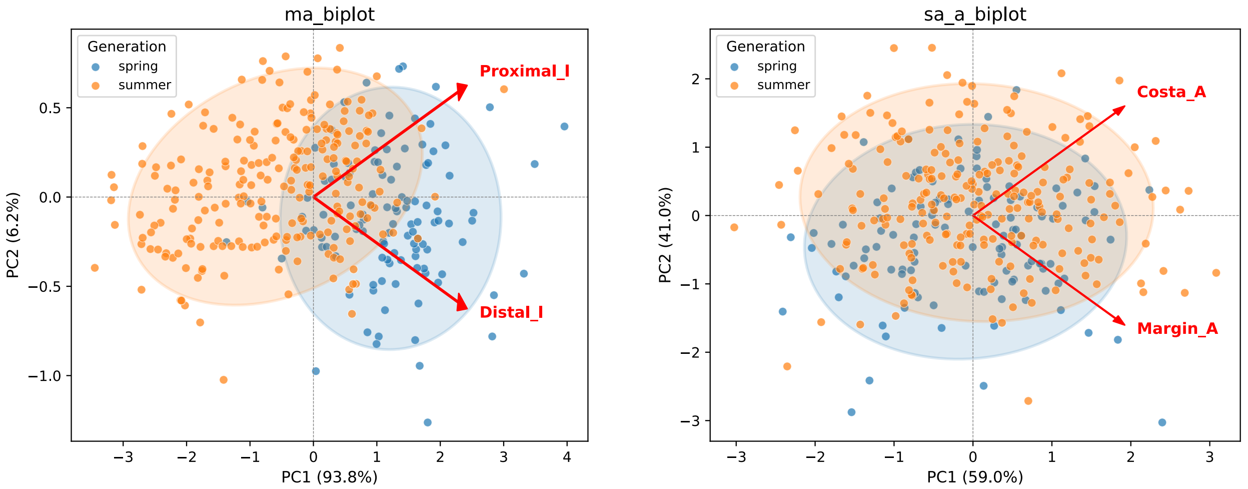


Figure 1: Biplot of the proximal and distal segments of the wing longitudinal axis (left) and the costal and marginal angles (right). PC1 captures overall size; PC2 captures the inverse relationship between the two segments/angles, corresponding to the MA and SA metrics, respectively.

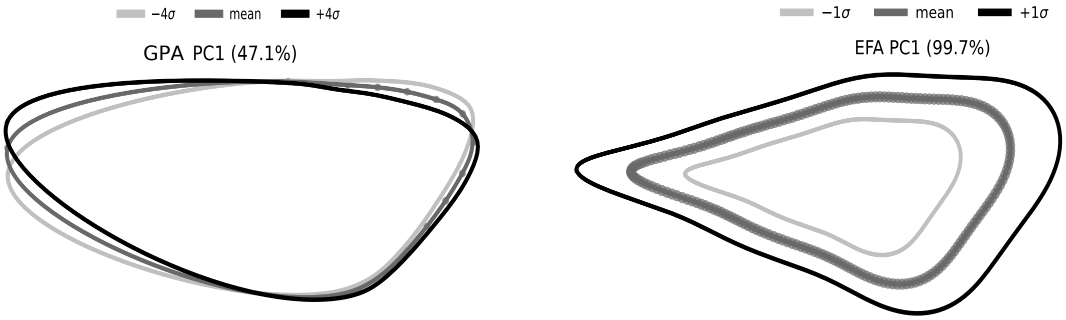


Figure 2: Visualization of the shift error (random error) (left) and the size component (right) and their contribution to data variation. (left) Small displacements of the anchor landmark shift the entire landmark chain. (right) The first EFA principal component on raw (unaligned) landmarks corresponds to overall size.
